# Identification of novel and structurally diverse N-Methyl-D-Aspartate Receptor Antagonists: Successful Application of Pharmacophore Modeling, Virtual Screening and Molecular Docking

**DOI:** 10.1101/314914

**Authors:** Mukta Sharma, Anupama Mittal, Aarti Singh, Ashwin K. Jainarayanan, Sarvesh Kumar Paliwal

## Abstract

In view of “excitotoxic” effects of glutamate, wherein excessive excitatory input causes increase in intracellular Ca^2+^ and ultimately cell death, NMDA receptor has emerged as an important target for treatment and prevention of several neurological disorders, like Alzheimer disease. Prompted by the successful application of *in-silico* pharmacophore-based virtual screening in lead identification, we have made an effort to implement *in-silico* protocols to identify novel NMDA receptor antagonist. A series of novel benzo[b]quinolizinium cations as NMDA receptor antagonists have been used as a starting point to develop prognostic pharmacophore models. The most predictive pharmacophore model (hypothesis 1), consisting of four features, namely, one hydrogen bond acceptor, one hydrophobic and two ring aromatic, showed a correlation (*r*) of 0.89, root mean square of 0.259, and the cost difference of 43.01 bits between null and fixed cost. The model was thoroughly validated and subjected to a chemical database search, which lead to the identification of 400 hits from NCI and Maybridge databases which were checked for Lipinski’s violation and predictive potency.

This reduced the list to 10 compounds, out of which, two most potent compounds were subjected to molecular docking using Libdock software and interestingly, all the docked conformations showed hydrogen bond interactions with important amino acids Tyr214, His88, Thr174, Val169 and Arg121. In summary, through our validated pharmacophore-based virtual screening protocol, we have identified two potent, structurally diverse, druggable and novel NMDA receptor antagonist which might be of great help to address the unmet medical need of Alzheimer disease.

## 1. INTRODUCTION

Alzheimer’s disease (AD), a progressive neurodegenerative (Crews and Masliah, 2010) disease described by a decline in mental stability and neuronal loss is associated with three main pathological properties which include: occurrence of neurofibrillary tangles, senile plaques (Pozo et al., 2011) and loss of synapses. Alzheimer’s disease has affected nearly 15 million people and in the absence of any proper treatment, the number of patients is likely to increase to 22 million worldwide and it is noticeable that, AD is present in nearly half of individuals aged above 80 years. The neuronal damage observed in AD is considered to be an outcome of the NMDA receptor-mediated “excitotoxic” effects of glutamate (Danysz and Parsons, 2012) wherein excessive excitatory input causes an increase in intracellular Ca^2+^ and ultimately cell death. NMDARs mainly comprise a sub-family with definite molecular composition and unusual pharmacological and functional properties. These receptors play a significant role in both synaptic plasticity (Spalloni et al., 2013) and neuronal death. They are generally heteromeric complexes including diverse subunits (Awobuluyi et al., 2007) which include mainly three subtypes: NR1, NR2 and NR3. NR1 contains eight different subunits created by alternative splicing from a single gene, NR2 and NR3 contain four (A, B, C and D) and two (A and B) different subunits respectively. The NR1 subunit is comprised of a glycine binding site while the homologous domain on the NR2 subunit contains the glutamate (Dong et al., 2009) binding site. The importance of NMDARs in the pathogenesis of AD makes them an attractive target for the development of anti-Alzheimer drugs.

The role of NMDA receptor antagonist like memantine (Sonkusare et al., 2005) in prevention and treatment (Parsons et al., 1999) of neurodegenerative disorder has further strengthened the concept of use of such class of drugs in AD. In present work, we have disclosed our effort to identify novel and structurally diverse NMDA receptor antagonists through a well defined sequential *in-silico* virtual screening protocol.

## 2. MATERIALS AND METHODS

A series of 24 compounds belonging to the novel benzo[b] quinolizinium (Earley et al., 1995) cations antagonists along with their corresponding biological data represented as Ki values in nM were employed for the pharmacophore generation. All the molecules under consideration were randomly split into training and test set comprising of 17 and 07 compounds respectively. Energy minimization was carried for all molecules under study using CHARMm force field and conformations (Peng et al., 2010) were generated using the ‘‘best’’ option with an energy cut-off of 20 kcal/mol.

### 2.1 Generation of pharmacophore

All molecules in the training set were used for hypothesis (pharmacophore) generation employing hypogen module of discovery studio (Accelrys, 2005). Hypogen aims to identify the best 3-dimensional arrangement of chemical functions explaining the activity variations among the compounds in the training set. Precisely HypoGen algorithm attempts to find hypotheses that are common among the active compounds of the training set. Instead of using just the lowest energy conformation of each compound, all the conformational models for molecules in each training set were used for pharmacophore hypothesis generation. During the hypothesis generation, it was observed that four features, i.e., one hydrogen bond acceptor (HBA), one hydrophobic (HY) and two ring aromatic (RA) features, dominated in most of the useful hypotheses generated. Therefore, these four features were used to construct a final set of 10 pharmacophore hypotheses from the training set, using a default uncertainty value of 3.

The quality of the generated pharmacophore models were evaluated on the basis of correlation coefficient, RMS and cost function analysis. The cost value (total cost) consists of three components namely, the weight cost, the error cost and the configuration cost (Arooj et al., 2011).

The weight component increases in a Gaussian form as the feature weight deviates from the ideal value of 2.0. The error cost increases as the RMS distance between the estimated and the measured activities for the training set increases and the configuration cost represents the complexity or the entropy of the hypothesis space being optimized and is constant for a given data set. It depends on the complexity of the pharmacophore hypothesis space. Any value higher than 17 may indicate that the correlation from any of the generated hypothesis (Tsai et al., 2006) is most likely due to chance, so either some attention has to be given in the selection of training set molecules or the entropy cost should be reduced by limiting the minimum and maximum features. The generated pharmacophore models were also evaluated on the basis of two another important costs that are fixed cost and the null cost, the former represents the simplest model that perfectly fits the data and the null cost is the cost of a pharmacophore without any feature where the calculated activity data of each molecule in the training set is the average value of all activities. The differences between the cost of the generated hypothesis and the null hypothesis should be as large as possible; a value of 40–60 bits difference may indicate that there is 75–90% chance of representing a true correlation in the data set used. The total cost of any hypothesis should be nearer to the value of the fixed cost for any meaningful model. The RMS deviation represents the quality of the correlations between the estimated and the actual data.

### 2.2 Cat scramble validation

During Cat Scramble the biological data and the corresponding structures are scrambled several times and the pharmacophoric models are generated from the randomized data. The confidence in the parent hypotheses (i.e., generated from unscrambled data) is lowered proportional to the number of times the software succeeds in generating binding hypotheses from scrambled data of apparently better cost criteria than the parent hypotheses. The statistical significance is given by the equation.

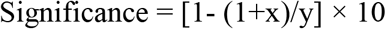

Where x = total number of hypotheses having a total cost lower than best significant hypothesis and y = number (HypoGen runs initial+ random runs).

The best pharmacophore hypothesis is chosen on the basis of the correlation coefficient, RMSD and cost function analysis results were evaluated for its statistical significance using the aforesaid Cat Scramble program. To obtain a 95% confidence level, 19 random spreadsheets were generated (y= 20) and every generated spreadsheet was submitted to HypoGen using the same experimental conditions (functions and parameters) as the initial run and the results were analyzed.

### 2.3 Internal test set prediction

An internal test set comprising 07 compounds was employed to assess the prediction ability of the developed model. All the selected test set compounds were mapped onto the generated pharmacophoric model and thus the prediction of the desired activity was made. The Catalyst program fits each compound to a hypothesis and reports back a series of ‘Fit’ scores. The fit function does not depend only on the mapping of the feature but also possesses a distance term measuring the distance between the feature on the molecule and the centroid of the hypothesis feature, and both these terms are used in the calculation of geometric fitness. Squared correlation coefficient value between actual activities and the corresponding predicted activities for all test set molecules was observed after mapping of molecules onto the hypothesis.

### 2.4 External test set validation

As a final step towards validation, the predictive power of the chosen pharmacophoric model was judged using an external test set comprising structurally different NMDA receptor antagonists. All the test set molecules were built and treated analogous to training set compounds and mapped onto the HypoGen pharmacophore. The mapping pattern fit and estimated values were critically analyzed for all the compounds under consideration.

### 2.5 Database screening

It is an established fact that the pharmacophore model, if validated properly, provides a precise idea of the required molecular features for a new lead and hence can be used as a search query to retrieve molecules with novel and desired attributes from chemical compound databases. Thus, with the purpose of identifying the novel, potent NMDA receptor antagonists, the four-feature pharmacophore model was used as a query to search NCI and Maybridge databases.

The retrieved hits were subjected to rigorous filtering on the basis of the fit score and Lipinski’s violation (Lipinski, 2000).

### 2.6 Molecular docking

In order to elucidate the binding interaction between NMDA receptor and the hits retrieved from the virtual screening, the molecular docking studies were performed using LibDock program which is an extension of DS (2.0). The crystal structure of NMDA [PDB ID: 3OEK] in complex with NAG was obtained from the Protein Data Bank (Vance et al., 2011) and was checked for valency, missing hydrogens and any other structural disorders like connectivity, bond orders, etc. The selected protein was split into the protein and ligand part. The protein was defined as receptor molecule whereas crystal ligand was used to define the binding site of 9 Angstroms on the receptor molecule. The structures of identified hits were sketched and energy minimized to obtain the most stable structure for docking. Memantine (Johnson and Kotermanski, 2006), which is FDA approved drug, was used as a reference for docking analysis. Based on coordinates, most potent hits and memantine were docked into the active site of NMDA receptor. Finally, all the possible interaction modes for different alignments were analyzed on the basis of Libdock (Rao et al., 2007) Scores.

## 3. RESULT AND DISCUSSION

### 3.1 Pharmacophore generation

The HypoGen algorithm was applied on the training set of 17 compounds with NMDA inhibitory activity (Table 1) to generate pharmacophore hypotheses. One compound (26) showed a greater error ratio, so it was considered as a potential outlier, hence removed during the course of pharmacophore modeling. The quality of the generated pharmacophore hypotheses was evaluated by considering the cost functions represented in bits unit calculated by HypoGen module during pharmacophore generation. The fixed cost of the 10 top-scored hypotheses was 72.26 bits, well separated from the null hypothesis cost of 115.27 bits. The total hypothesis cost, expressed in bits, of the 10 best hypotheses varied from 87.8 to 100.2. Such a range, covering only 13 bits is always desirable for healthy pharmacophore model building. The cost values, correlation coefficients (r), RMSD, and features for the top ten hypotheses are listed in Table 2.

**Table 1:**
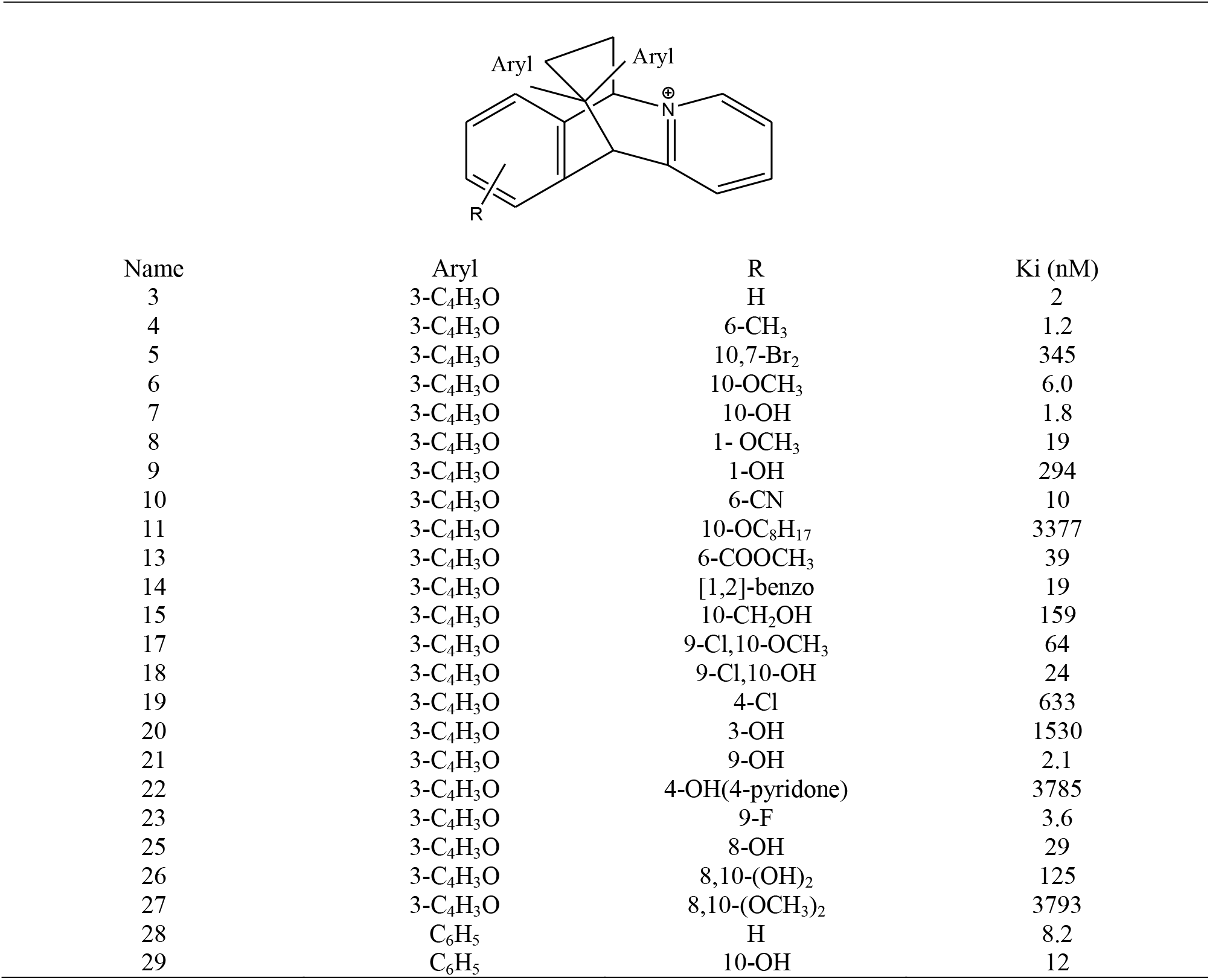
The molecular structures of the novel benzo[b]quinolizinium cations and their Ki values taken from the literature

|  |  |  |  |
| --- | --- | --- | --- |
| Name | Aryl | R | Ki (nM) |
| 3 | 3-C<sub>4</sub>H<sub>3</sub>O | H | 2 |
| 4 | 3-C<sub>4</sub>H<sub>3</sub>O | 6-CH<sub>3</sub> | 1.2 |
| 5 | 3-C<sub>4</sub>H<sub>3</sub>O | 10,7-Br<sub>2</sub> | 345 |
| 6 | 3-C<sub>4</sub>H<sub>3</sub>O | 10-OCH<sub>3</sub> | 6.0 |
| 7 | 3-C<sub>4</sub>H<sub>3</sub>O | 10-OH | 1.8 |
| 8 | 3-C<sub>4</sub>H<sub>3</sub>O | 1-OCH<sub>3</sub> | 19 |
| 9 | 3-C<sub>4</sub>H<sub>3</sub>O | 1-OH | 294 |
| 10 | 3-C<sub>4</sub>H<sub>3</sub>O | 6-CN | 10 |
| 11 | 3-C<sub>4</sub>H<sub>3</sub>O | 10-OC<sub>8</sub>H<sub>17</sub> | 3377 |
| 13 | 3-C<sub>4</sub>H<sub>3</sub>O | 6-COOCH<sub>3</sub> | 39 |
| 14 | 3-C<sub>4</sub>H<sub>3</sub>O | [1,2]-benzo | 19 |
| 15 | 3-C<sub>4</sub>H<sub>3</sub>O | 10-CH<sub>2</sub>OH | 159 |
| 17 | 3-C<sub>4</sub>H<sub>3</sub>O | 9-Cl,10-OCH<sub>3</sub> | 64 |
| 18 | 3-C<sub>4</sub>H<sub>3</sub>O | 9-Cl,10-OH | 24 |
| 19 | 3-C<sub>4</sub>H<sub>3</sub>O | 4-Cl | 633 |
| 20 | 3-C<sub>4</sub>H<sub>3</sub>O | 3-OH | 1530 |
| 21 | 3-C<sub>4</sub>H<sub>3</sub>O | 9-OH | 2.1 |
| 22 | 3-C<sub>4</sub>H<sub>3</sub>O | 4-OH(4-pyridone) | 3785 |
| 23 | 3-C<sub>4</sub>H<sub>3</sub>O | 9-F | 3.6 |
| 25 | 3-C<sub>4</sub>H<sub>3</sub>O | 8-OH | 29 |
| 26 | 3-C<sub>4</sub>H<sub>3</sub>O | 8,10-(OH)<sub>2</sub> | 125 |
| 27 | 3-C<sub>4</sub>H<sub>3</sub>O | 8,10-(OCH<sub>3</sub>)<sub>2</sub> | 3793 |
| 28 | C<sub>6</sub>H<sub>5</sub> | H | 8.2 |
| 29 | C<sub>6</sub>H<sub>5</sub> | 10-OH | 12 |

**Table 2:** Results of the top 10 hypotheses.

| Hypotheses | Total cost | Cost difference | RMSD | Correlation | Feature |
| --- | --- | --- | --- | --- | --- |
| 1 | 87.81 | 27.46 | 0.25 | 0.89 | HBA,HY,2RA |
| 2 | 89.18 | 26.09 | 0.48 | 0.84 | HBA,HY,2RA |
| 3 | 91.89 | 23.38 | 0.90 | 0.83 | HBA,HY,2RA |
| 4 | 91.90 | 23.37 | 1.21 | 0.82 | HBA,HY,2RA |
| 5 | 93.58 | 21.69 | 0.89 | 0.80 | HBA,HY,2RA |
| **6** | 94.80 | 20.47 | 0.45 | 0.78 | HBA,HY,2RA |
| **7** | 97.21 | 18.06 | 1.81 | 0.80 | HBA,HY,2RA |
| **8** | 98.90 | 16.37 | 1.20 | 0.85 | HBA,HY,2RA |
| **9** | 99.12 | 16.15 | 0.40 | 0.84 | HBA,HY,2RA |
| **10** | 100.2 | 15.07 | 1.92 | 0.72 | HBA,HY,2RA |

The top-ranked pharmacophore model (hypo 1) with one hydrogen bond acceptor (HBA), one hydrophobic (HY) and two ring aromatic features showed a high squared correlation coefficient of 0.89, low root-mean-square deviation of 0.25, weight of 1.29 and cost difference of 27.46, satisfying the acceptable range recommended in the cost analysis of the Catalyst procedure as shown in Fig. 1 (a). The configuration cost was 14.23, which is in agreement to standard requirement of configuration (Ashton et al., 1994) cost that should not exceed a maximum value of 17 (corresponds to a number of 2^17^ pharmacophore models) because high values may lead to a chance correlation of the generated hypothesis. Thus, hypothesis 1 was considered as the best pharmacophore model for NMDA inhibitory activity and was used to predict the activity of all the 17 training set compounds. A plot between the observed versus estimated activity demonstrated a good correlation coefficient (r^2^ training = 0.79) for training set compounds, indicating the high predictive ability of the pharmacophore as shown in Fig. 1 (b). The most potent NMDAR antagonist, **4**, mapped all features of the best pharmacophore model with the least displacement from the centroid, on the other hand the, least potent compound **27** missed to map one feature that is one HBA, (Fig. 1 (c), (d)).

**Fig. 1.**
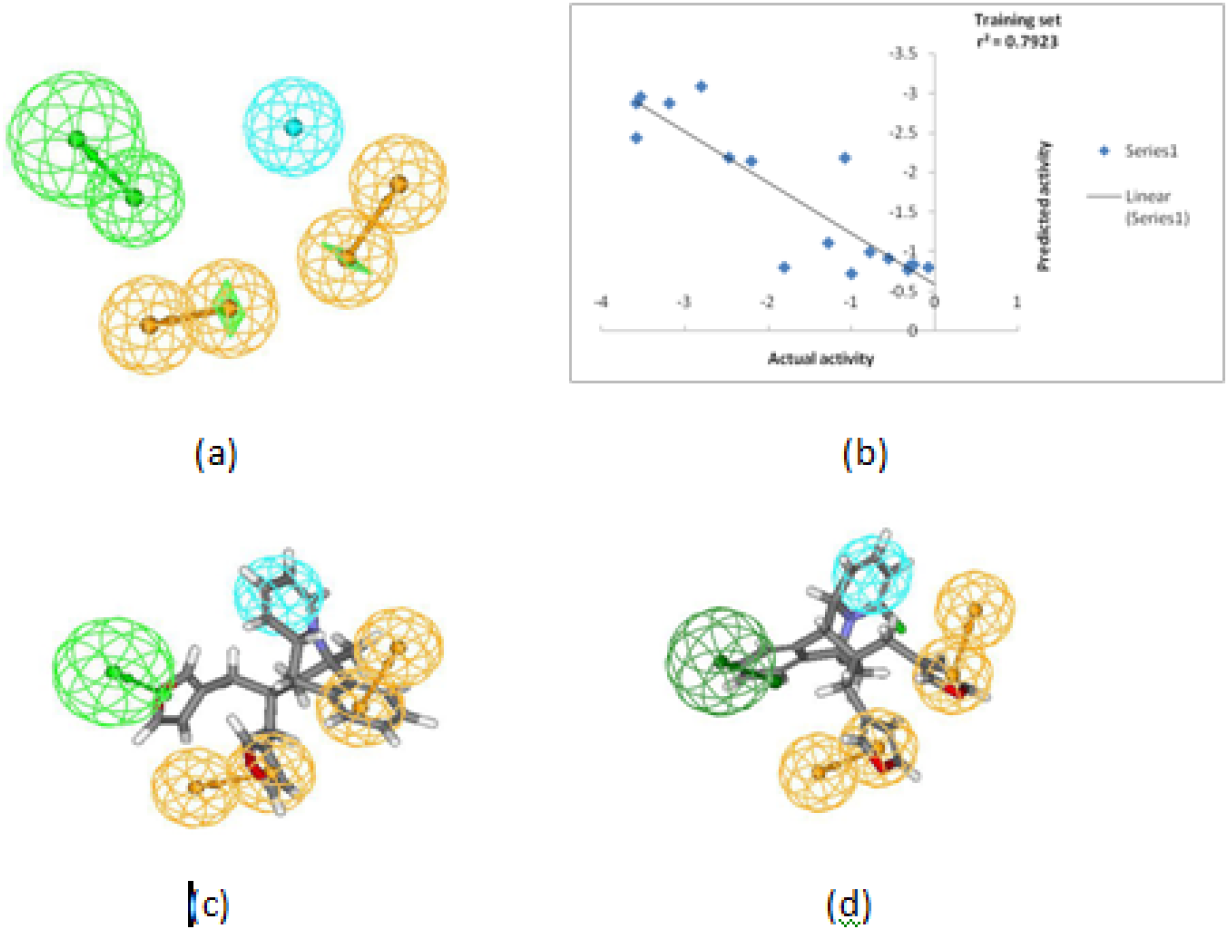
(a) Pharmacophoric features identified from best hypothesis 1. (b) A plot of actual versus estimated biological activity for training set compounds. (c) Mapping of most active compound 4 (Novel Benzo[b]quinolizinium derivative onto the chosen pharmacophore model (hypothesis 1). (d) Mapping of least active compound 27 (Novel Benzo[b]quinolizinium derivative) onto the chosen pharmacophore model (hypothesis 1)

### 3.2 Pharmacophore model validation

Once the chosen pharmacophore model satisfied the statistical criteria and successfully cleared the cost function test, it was rigorously validated using *Cat Scramble, internal and external test set prediction.*

#### (i) Cat scramble test

To evaluate the statistical relevance of the model, the Fischer validation method at the confidence level of 95% was applied to the developed HypoGen model and thus 19 spreadsheets were generated, Fig. 2. These random spreadsheets were used to generate hypotheses employing exactly the same features as used in generating the initial hypothesis. The experimental activities in the training set were scrambled randomly using Cat Scramble (Pal and Paliwal, 2012) program, and the resulting training set was used for a HypoGen run. In this manner, all parameters were taken from the initial HypoGen calculation. None of the outcome hypotheses had a better statistics than the initial hypothesis which verified that the hypothesis 1 has not been obtained by chance, Fig. 3. This, validation method provided enough confidence in the chosen pharmacophore model.

**Fig. 2.**
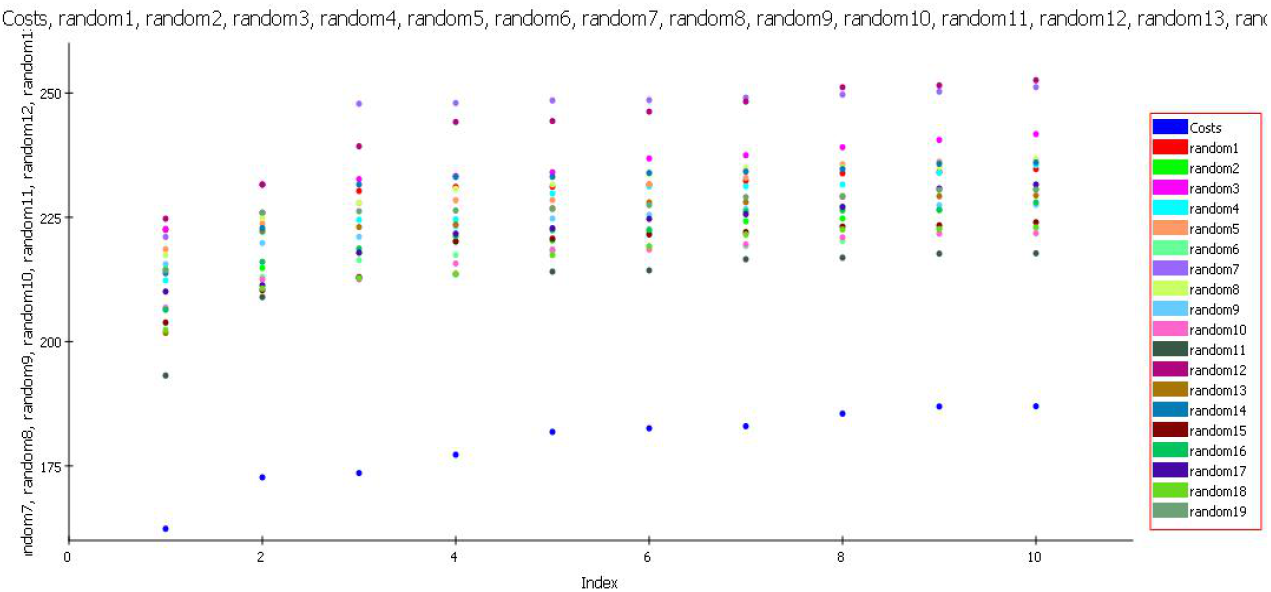
Graph of 95% cat scrambled cost data. None of the outcome hypotheses had a higher correlation score than the initial (best) hypothesis.

**Fig. 3.**
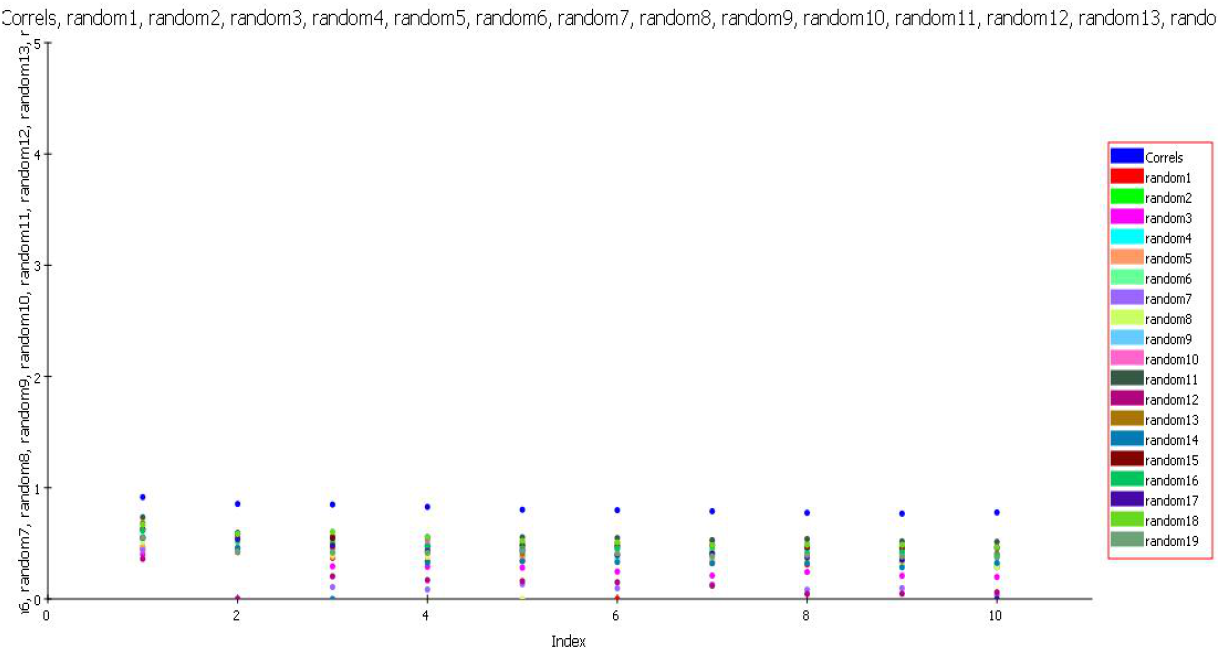
Graph of 95% cat scrambled correlation data. None of the outcome hypotheses had a higher correlation score than the initial (best) hypothesis.

#### (ii) Internal test set

The purpose of the pharmacophore hypothesis generation is not just to predict the activity of the training set compounds accurately but also to verify whether the pharmacophore model is capable of predicting the activities of compounds not included in the training set. A test set consisting of 07 ligands was subjected to pharmacophore mapping analysis employing the selected pharmacophore model. All molecules in the test set were built, minimized and subjected to conformational analysis like the molecules in the training set. Finally. the compounds were mapped onto the pharmacophore model using the best fit option. The results as shown in Fig. 4 clearly indicate towards a good correlation between the actual and estimated activities.

**Fig. 4.**
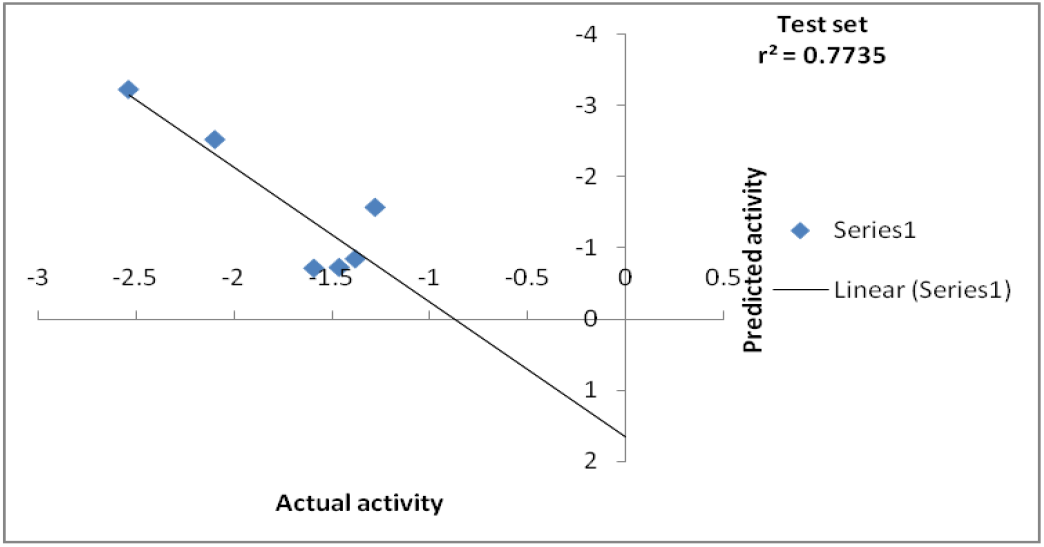
A plot of actual versus estimated biological activity for test set compounds

#### (iii) External test set validation

Finally, the predictive power of the selected pharmacophore model was evaluated with the help of an external test set comprising 10 NMDAR antagonists with high level of structural diversity. All the compounds were mapped onto the developed pharmacophore model and interestingly all the molecules of external test set exhibited a perfect four–feature mapping with good fit values. Squared correlation coefficient value of 0.82 between actual and estimated value of external test set compounds confirmed that the pharmacophore model has a high level of predictive power and universality, Fig. 5. The mapping of the best fit molecule of the external test set is shown in Fig. 6.

**Fig. 5.**
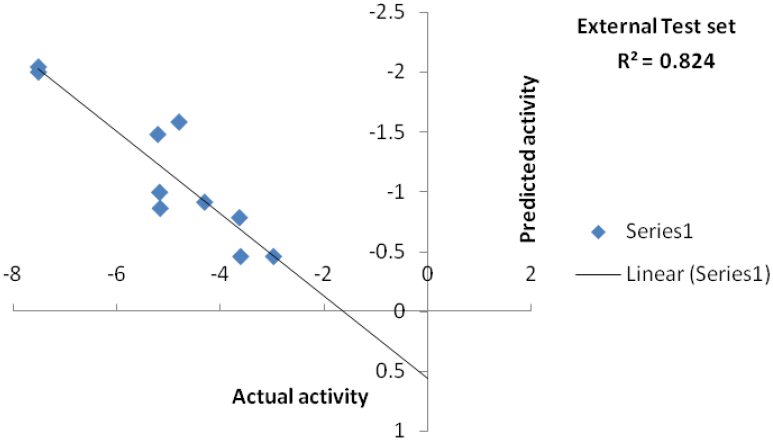
A plot of actual versus estimated biological activity for external test set compounds.

**Fig. 6.**
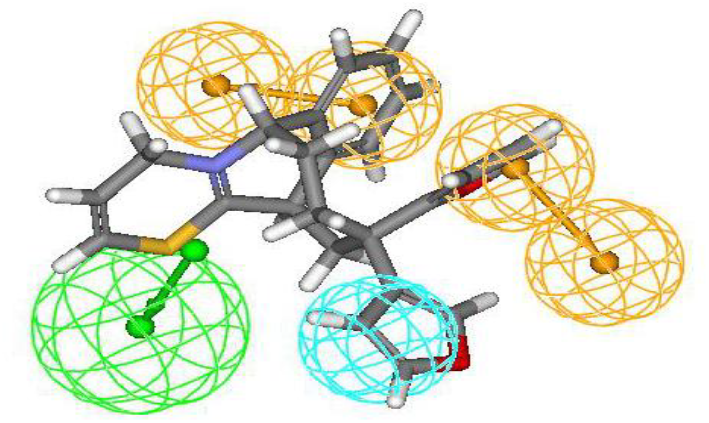
Pharmacophore mapping of most active compound of external test set onto the chosen pharmacophore model (hypothesis 1).

### 3.3 Database screening

The validated pharmacophore model was used as a query for virtual screening of NCI and Maybridge chemical compound databases to identify novel, potential lead compounds with good fit values. Database screening (Langer and Krovat, 2003) led to the retrieval of 400 hits. All the hits were checked for their fit values and Lipinski’s violation, reducing the list to ten compounds namely NSC3665, NSC2402, NSC3178, NSC390, NSC1012, JFD03984, HTS03850, BTB08221, BTB14180, HTS04999 which turned out as potential ligands exhibiting a perfect four feature mapping with good fit value ranging from 7.452-6.329 and zero Lipinski’s violation, Table 3. Out of ten, two most potent hits were subjected to molecular docking to observe the molecular interactions of the selected hits.

**Table 3:**
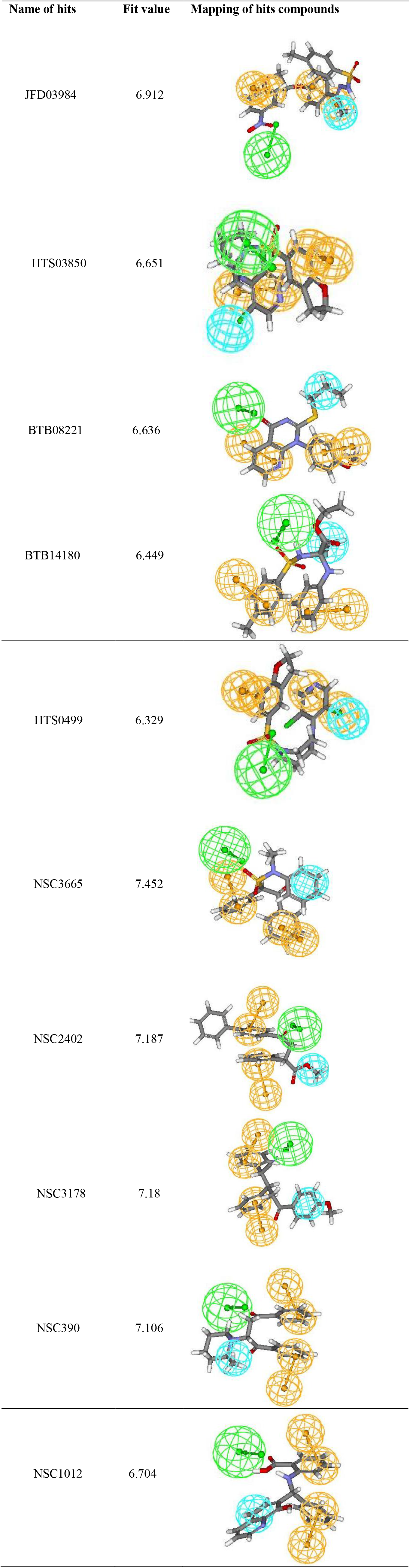
Hits retrieved from Maybridge and NCI databases.

| Name of hits | Fit value | Mapping of hits compounds |
| --- | --- | --- |
| JFD03984 | 6.912 |  |
| HTS03850 | 6.651 |  |
| BTB08221 | 6.636 |  |
| BTB14180 | 6.449 |  |

### 3.4 Molecular docking

NMDA receptor [PDB ID: 3OEK] in complex with NAG (N-Acetyl-Glucosamine) was obtained from the Protein Data Bank for molecular docking studies. The structures of NSC2402 (NCI), HTS03850 (Maybridge) and memantine were prepared and docked in the active site of 3OEK using LibDock software which is a molecular dynamics simulated annealing based algorithm (Meng et al., 2011) and is available as an extension of DS V.2.0. Total 100 poses were generated for NSC2402, HTS03850 and memantine and all the poses were inspected for the type of interaction between chosen compounds and binding site of NMDA receptor. It was observed that NSC2402 and HTS03850 exhibited high LibDock score of 116.651 and 153.539 respectively and interestingly the drug memantine showed a comparatively low LibDock score of 70.16.

The interaction analysis of NSC2402 showed that oxygen present on the propyl-benzene ring is interacting with Tyr214. Hydrogen present on the 2^nd^ position of toluene ring exhibited Vander Waals interactions with Val169 and Tyr214. On the other hand hydrogen present on the 3^rd^ position of toluene ring showed Vander Waals interactions with Thr174 and hydrogen present on methyl acetate group showed Vander Waals interactions with His88 as shown in Fig. 7. In case of HTS03850 dimethylamine group formed hydrogen bond with Val169. Nitrogen of dimethylamine group and hydrogen present on the 2^nd^ position of benzodioxole ring showed Vander Waals interaction with Tyr214. Oxygen present on the 3^rd^ position of 1, 3-dioxolane ring showed hydrogen bond interaction with Arg121.Hydrogen present on dioxolane ring showed Vander Waals interactions with Arg121 and hydrogen present on the 6^th^ position of benzodioxole ring showed Vander Waals interaction with Gly172 as shown in Fig. 8.

**Fig. 7.**
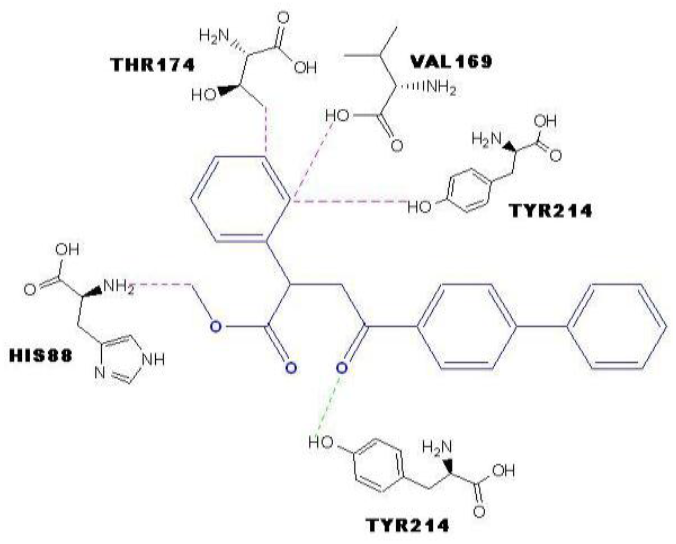
Interaction of NSC2402 with Tyr214, Val169, Tyr214, Thr174, His88, the active site amino acids. [Green dotted lines representing Hydrogen bond interactions; Pink dotted lines representing Vander Waals interaction]

**Fig. 8.**
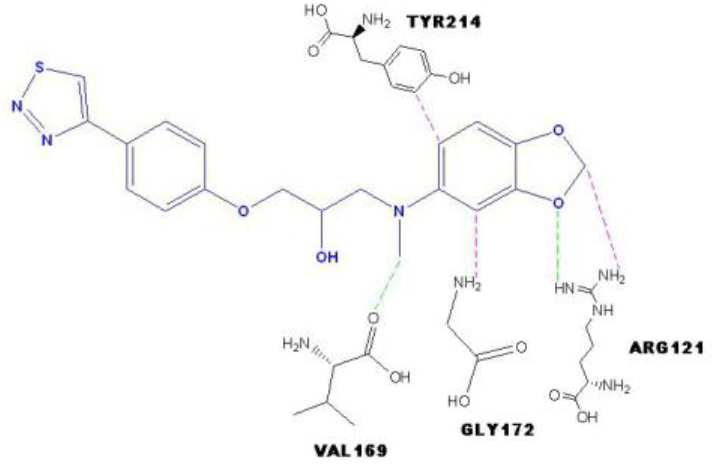
Interaction of HTS03850 with Val169, Tyr214, Arg121, Gly172 the active site amino acids. [Green dotted lines representing Hydrogen bond interactions; Pink dotted lines representing Vander Waals interaction]

While comparing the docking results of two lead molecules, it was observed that hydrogen bond and Vander Waal interactions with essential amino acids are contributing heavily towards the activity of the NSC2402 and HTS03850. Interaction analysis of reference drug memantine revealed that amine group present on adamantan-1-amine ring showed hydrogen bond interactions with Ser131, Tyr282, Gly264, Ser260 and Ser26. Methyl present on the 3^rd^ and 5^th^ position of amine ring showed Vander Waals interactions with Asp283, His127 and Arg292 as shown in Fig. 9.

**Fig. 9.**
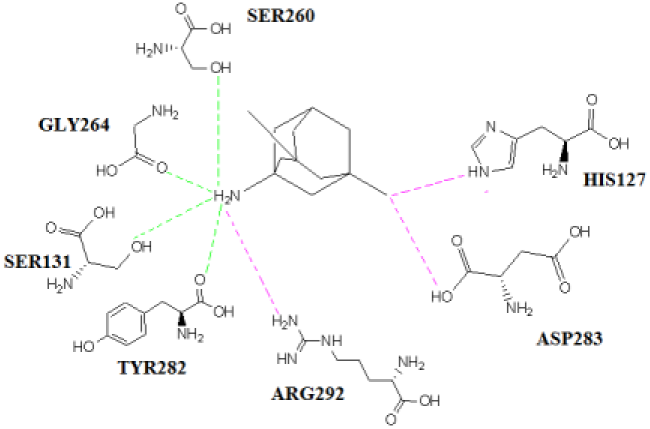
Interaction of Memantine with Ser131, Tyr282, Gly264, Ser260, Ser26, Asp283, His127, Arg292 the active site amino acids. [Green dotted lines representing Hydrogen bond interactions; Pink dotted lines representing Vander Waals interaction]

### 3.5 Novelty check of NSC2402 and HTS03850

In view of good fit value, drug-likeness and docking score, NSC2402 and HTS03850 were checked for novelty by employing pairwise Tanimoto (Shen et al., 2013) similarity indices using “Find Similar Molecules by Fingerprint” protocol in Discovery Studio. Both the compounds showed low Tanimoto similarity indices of 0.169 and 0.077 to all the structures of known NMDA Receptor antagonists confirming their novelty. Since the number of AD patient is increasing day by day and there is a definite scarcity of efficacious therapeutic options, the identified novel and potent NMDA receptor antagonists could be sound template for the development of new anti-Alzheimer drugs.

## 4. CONCLUSION

In view of the unmet medical need of Alzheimers Disease, we have developed a rigorously validated pharmacophore model for NMDA receptor antagonists. The generated pharmacophore reflects the binding mode and the important interactions of the ligands with amino acids in the active site of NMDA. The best pharmacophore model consisting of one hydrogen bond acceptor (HBA), one hydrophobic (HY) and two ring aromatic features satisfied all the statistical criteria. The model was judiciously used to identify novel hits through database mining. This led to the identification of two most potent compounds which were subjected to molecular docking studies. Both the compounds showed good LibDock scores and interaction with important active site amino acid residues (Tyr214, Val169, and Arg121). In summary, through our reported research endeavor, we have identified novel, potent and druggable candidates which could be raised into potential neuroprotective agents.

## ACKNOWLEDGMENT

We are grateful to Prof. Aditya Shastri, Vice Chancellor, Banasthali University (India) for providing necessary computational facilities for the study.

## AUTHOR CONTRIBUTIONS

MS performed the whole simulation experiment. SKP has made major contributions to conception and design of the experiments. AM, AS and AKJ supported in analyzing the results. MS wrote the manuscript.

